# Intra-saccadic motion streaks jump-start gaze correction

**DOI:** 10.1101/2020.04.30.070094

**Authors:** Richard Schweitzer, Martin Rolfs

## Abstract

Rapid eye movements (saccades) incessantly shift objects across the retina. To establish object correspondence, the visual system is thought to match surface features of objects across saccades. Here we show that an object’s intra-saccadic retinal trace – a signal previously considered unavailable to visual processing – facilitates this match-making. Human observers made saccades to a cued target in a circular stimulus array. Using high-speed visual projection, we swiftly rotated this array during the eyes’ flight, displaying continuous intra-saccadic target motion. Observers’ saccades landed between the target and a distractor, prompting secondary saccades. Independently of the availability of object features, which we controlled tightly, target motion increased the rate and reduced the latency of gaze-correcting saccades to the initial pre-saccadic target, in particular when the target’s stimulus features incidentally gave rise to efficient motion streaks. These results suggest that intra-saccadic visual information informs the establishment of object correspondence and jump-starts gaze correction.

## Introduction

Saccadic eye movements are the fastest and most frequent movements of the human body. By placing the fovea at new parts of the visual scene, they provide high-acuity vision across the visual field. At the same time, saccades result in retinal translations that constantly shift the projections of objects onto the retina, and impose significant motion blur on the retinal image. These consequences, however, do not impair our visual experience, a phenomenon widely known as visual stability (Bridgeman et al., 1994; MacKay, 1973; McConkie & Currie, 1996; Wurtz, 2008). A core component of visual stability is the establishment of object correspondence across saccades: How does the visual system determine whether any object located in the periphery prior to a saccade is the same as the object close to the fovea right after that saccade has landed?

There is good evidence that visual short-term memory (VSTM) enables the matching of objects across saccades (Aagten-Murphy & Bays, 2019). For instance, Hollingworth et al. (2008) showed that surface features of visual objects encoded in VSTM, such as color or object identity, can be used for gaze correction when targets were displaced during saccades. To some extent, this result contradicted object-file theory (Kahneman et al., 1992), which supports the notion that objects are referenced via spatiotemporal continuity, not surface features (Mitroff & Alvarez, 2007). Later studies then suggested that both surface features and spatiotemporal continuity could contribute to object correspondence, across both brief occlusions while fixating (Hollingworth & Franconeri, 2009) and across saccades (Richard et al., 2008).

One potential source of information for object correspondence has been neglected by all studies up to this point: Intra-saccadic object motion across the retina may provide spatiotemporal continuity as well as access to surface features before saccade landing. As illustrated in **Figure 1**, due to temporal integration in the visual system, objects moving at the high velocities of saccades routinely produce smeared traces, so-called motion streaks (Bedell & Yang, 2001; Brooks et al., 1980; Duyck et al., 2016; Geisler, 1999; Matin et al., 1972). Most experiments thus far, in fact, were built on the premise that “vision is suppressed, creating a gap in perceptual input” (Richard et al., 2008, p. 66) and “people are virtually blind” (Hollingworth et al., 2008, p. 163) during saccades. In contrast to this premise, we have recently shown that observers can use intra-saccadic motion streaks to tell pre-saccadic objects from identical distractors upon saccade landing (Schweitzer & Rolfs, 2020). The crucial question — if intra-saccadic streaks could be used to establish object correspondence across saccades — remains unanswered, however. To test this idea unequivocally, implicit behavioral measures rather than explicit perceptual reports must be used, as perceptual reports may draw observers’ attention to a source of information they might have otherwise ignored. Here, using a high-speed projection system, we adapted a classic gaze correction paradigm (Hollingworth et al., 2008) to investigate the hypothesis that continuous object motion — exclusively present during saccades — may serve object continuity and, hence, facilitate gaze correction.

**Figure 1.**
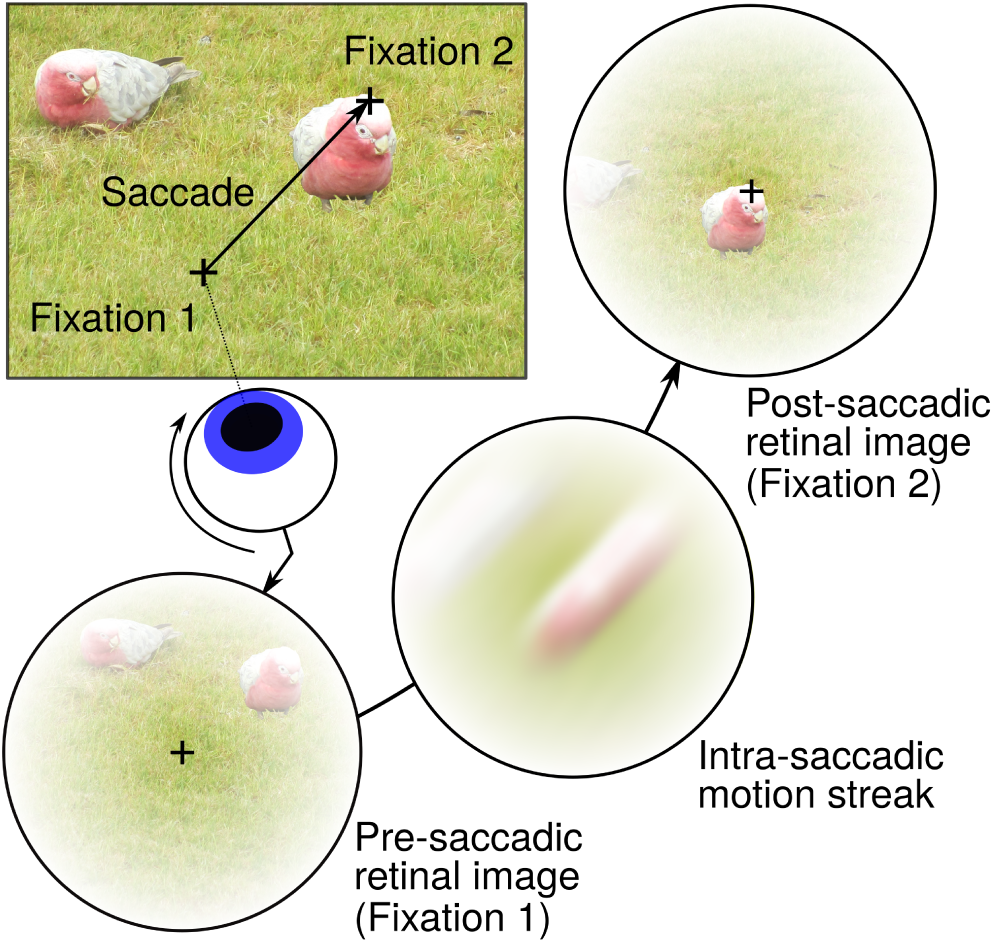
Illustration of intra-saccadic motion streaks. When making a saccade towards the bird on the right, its retinal projection rapidly travels from a pheripheral location (Fixation 1) to a foveal location (Fixation 2), producing a motion streak along its retinal trajectory. This streak literally connects an object’s pre- and post-saccadic locations on the retina, possibly providing spatiotemporal continuity that may help establish object correspondence. Even though intra-saccadic motion streaks are largely omitted from conscious visual perception, likely due to masking by pre- and post-saccadic retinal images (Campbell & Wurtz, 1978; Matin et al., 1972) and attenuation by saccadic suppression (Ross et al., 2001; Volkmann, 1986), could they still inform trans-saccadic processes?

## Methods

### Apparatus

Stimuli were projected onto a 16:9 (250.2 x 141.0 cm) video-projection screen (Stewart Silver 5D Deluxe, Stewart Filmscreen, Torrance, CA), mounted on a wall 340 cm in front of the participant, using a PROPixx DLP projector (Vpixx Technologies, Saint-Bruno, QC, Canada) running at 1440 Hz refresh rate and a resolution of 960 x 540 pixels. The experimental code was implemented in MATLAB 2016b (Mathworks, Natick, MA, USA) on Ubuntu 18.04, using Psychtoolbox (Kleiner et al., 2007; Pelli, 1997), and was run on a Dell Precision T7810 Workstation supplied with a Nvidia GTX 1070 graphics card. Eye movements of both eyes were recorded via a TRACKPixx3 tabletop system (Vpixx Technologies, Saint-Bruno, QC, Canada) running firmware version 11 and at a sampling rate of 2000 Hz, while participants rested their head on a chin rest. A custom wrapper function toolbox was used to control the eye tracker, which is made publicly available on Github: https://github.com/richardschweitzer/TrackPixxToolbox.

### Participants

Ten observers completed three sessions with a duration of approximately 1 hour each on separate days and received 26 Euros as remuneration (plus 2 Euros for every 15 minutes of overtime). We obtained written informed consent from all participants prior to inclusion in the study. The study was conducted in agreement with the latest version of the Declaration of Helsinki (2013) and was approved by the Ethics board of the Department of Psychology at Humboldt-Universität zu Berlin. All observers (5 male, 5 female; mean age: 28; age range: 20 – 37) had normal or corrected-to-normal vision (20/20 ft acuity in the Snellen test; 4 observers wore glasses and 1 observer wore contact lenses). Seven of ten observers had right ocular dominance (established by a variant of the Porta test) and ten observers were right-handed. The experiment was pre-registered at Open Science Framework (OSF). In accordance with pre-registered exclusion criteria, four invited participants had to be replaced because they did not complete all three sessions. Pre-registration, data, and analysis scripts can be found at https://osf.io/aqkzh/.

### Procedure & Task

A six-stimulus circular array at an eccentricity of 10 degrees of visual angle (dva) was displayed while observers fixated an area with a 1.5 dva radius around a central fixation dot for 400 ms. The stimulus array contained two types of dissimilar noise patches (see Stimuli), in alternating order (**Figure 2a**, top row). Specific stimulus positions were at 0, 60, 120, 180, 240, and 300 (as shown in **Figure 2a**), or alternatively at 30, 90, 150, 210, 270, and 330 degrees relative to the central fixation dot (0 deg: below the fixation point). After successful fixation, an exogenous cue was presented to indicate the saccade target: The target stimulus – one of the six presented stimuli and one of the two types of noise patches – was enlarged linearly up to twice its initial size for 25 ms and then decreased for 25 ms until the initial size was restored. Saccades were detected online using the algorithm described by Schweitzer & Rolfs (2019) with parameters k=2, λ=10, and ϑ=40, on both eyes. As soon as the saccade was detected, the cued stimulus moved 30 deg in a clockwise (CW) or counterclockwise (CCW) direction for 14.6 ms – amounting to a distance traveled of 5.2 dva at a velocity of approximately 360 dva/s – or remained in its pre-saccadic location. Importantly, this 14.6-ms motion was either continuous (motion-present condition), i.e., presenting 21 frames of equally spaced stimulus positions along the circular trajectory (0.25 dva per frame), or apparent (motion-absent condition), i.e., presenting a blank screen between the first and final positions of the stimulus. In both motion conditions, all other noise patches were removed during this short and rapid stimulus motion. As soon as the moving stimulus reached its final position, all stimuli were displayed at their post-motion locations consistent with a 30-deg CW or CCW rotation of the stimulus array. Observers’ saccades thus landed between two dissimilar noise patches: One was always the target stimulus which was cued prior to saccade initiation, and the other one – the distractor – was an uncued and therefore irrelevant stimulus. As a consequence, a secondary saccade was made to the target (or erroneously to the distractor) in order to correct for the intra-saccadic displacement which occurred in two thirds of all trials. Crucially, a pixel noise mask (**Figure 2a**, bottom row) occluded the identity of all stimuli presented on the screen with varying delay relative to stimulus motion offset (0, 25, 50, 100, 200, 600 ms), thus limiting observers’ time to use stimulus surface features to guide their secondary saccades. This post-motion mask onset delay will henceforth be referred to as of surface-feature duration. 650 ms after stimulus motion offset, each trial was concluded.

**Figure 2.**
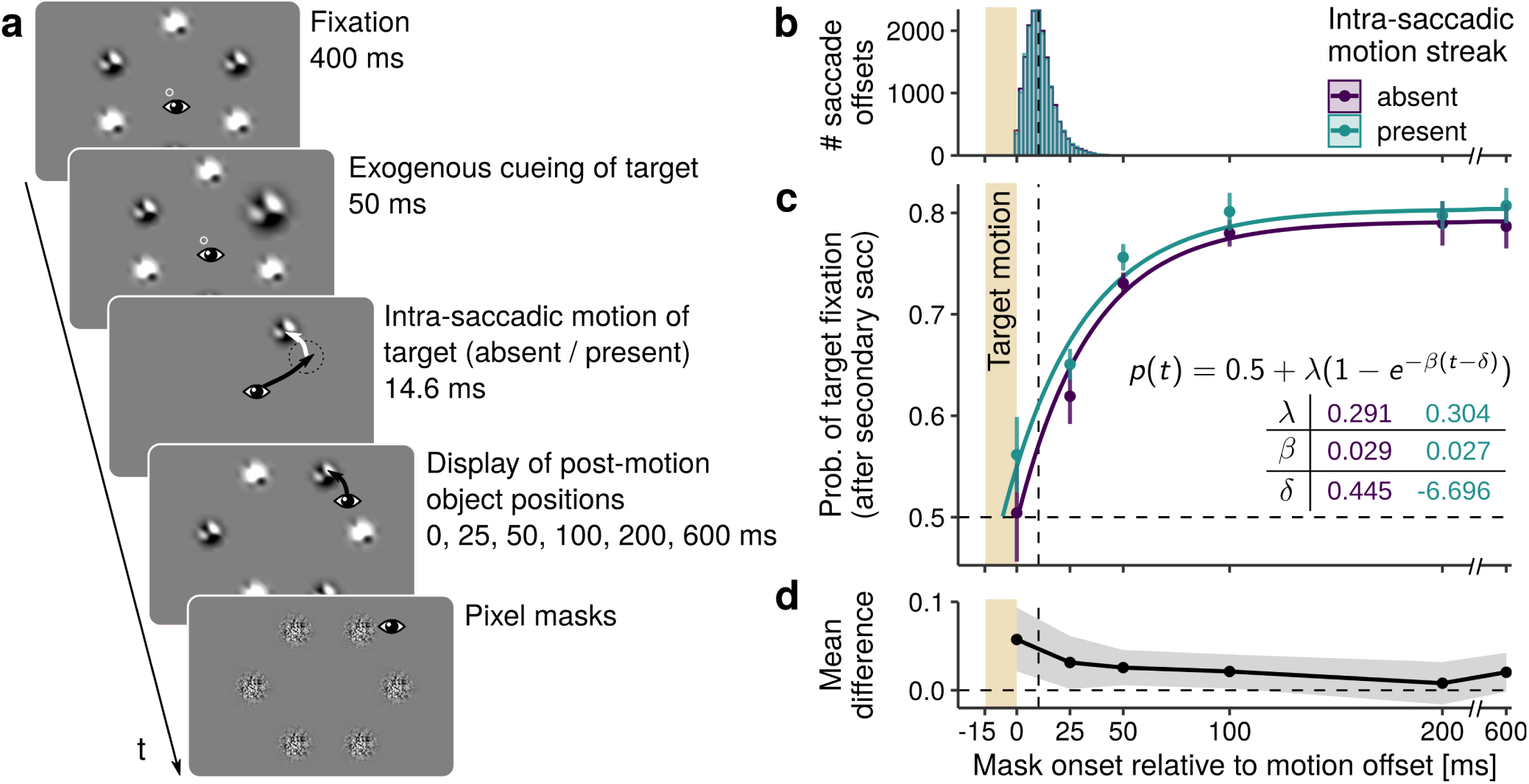
Probing the role of post-saccadic surface features and intra-saccadic motion in gaze correction. **a** Observers made a primary saccade to an exogenously cued target noise patch stimulus (one of two types). Strictly during the saccade, the target rapidly shifted positions, consistent with a 30-degree clockwise or counterclockwise rotation of the entire stimulus array, so that primary saccades always landed between the initially cued stimulus (the target) and the other-type stimulus (the distractor). The intra-saccadic stimulus motion was displayed for 14.6 ms and was either continuous (i.e., 21 equidistant steps along its circular trajectory) or absent (blank screen between first and final stimulus position). After the stimulus’ motion, pixel masks were displayed with a varying delay (surface-feature durations: 0, 25, 50, 100, 200, or 600 ms), thus occluding the identity of post-saccadic objects and limiting the observers’ ability to establish trans-saccadic correspondence using object features. **b** Stimulus motion was presented strictly during saccades, finishing on average 10.7 ms prior to saccade offset. **c** Likelihood of observers making a secondary saccade towards the initial pre-saccadic target was a function of surface-feature duration, as well as the presence of intra-saccadic motion (purple vs green points, respectively; error bars indicate ±*S EM*). The beige area illustrates the temporal interval in which intra-saccadic motion took place. Solid lines show predictions of the mixed-effects exponential growth model describing the increase of proportions with increasing surface-feature duration. Average parameter estimates are shown in the table below the model formula. **d** Mean differences between motion conditions for each surface-feature duration with corresponding 95% confidence intervals (gray-shaded area).

Similar to Hollingworth et al. (2008), observers were instructed to make a saccade to the target stimulus upon cue presentation and fixate it. They were informed that the stimulus array could rotate in some trials, in which case they could make a secondary saccade to follow the initial target. If observers’ initial central fixation was unsuccessful, or if their primary saccade did not end within a circular region of 2 dva around the pre-saccadic target location, or if more than one saccade was made to reach the pre-saccadic target location, appropriate feedback was provided, and the trial was repeated at the end of the session. No feedback related to observers’ secondary saccades was given. To elucidate the trial procedure, a 60-fps video (slowed down by a factor of 24 and using the mouse cursor as a representative of gaze position) can be found at OSF: https://osf.io/f48rm/.

### Stimuli

Stimuli were achromatic, random Gaussian noise patches (SD=1) bandpass-filtered to spatial frequencies (SF) from 0.25 to 1 cycles per dva (cpd), displayed on a uniform, grey background. One initial bandpass-filtered noise matrix was generated on each trial. To maximize the dissimilarity between the two types of noise patches, 75% of a noise SD was added to or substracted from the initial noise matrix, thus increasing or decreasing its luminance (for one example, see **Figure 2a**). This procedure inevitably led to some differences in spatial frequency and orientation for the two types of content in the pairs of noise patches. This effect was intended, as it allowed for both easier discrimination of the two types during trials and for reverse regression analyses involving stimulus features at a later stage (Schweitzer & Rolfs, 2020; Wyart et al., 2012). All noise patches were at full Michelson contrast to maximize their intra-saccadic visibility. Noise patches were enveloped in a Gaussian aperture with a standard deviation of 0.5 dva. Masks displayed at post-motion locations had the same dimensions, but consisted of random black-white pixel noise. Noise masks were identical copies for all stimuli.

The central fixation dot at the beginning of each trial consisted of a white circle of 0.3 dva radius. To indicate that the dot was fixated by the observer, the area within the circle was be filled by another white circle of 0.1 dva radius.

### Pre-processing

Observers completed at least 3456 trials, i.e., at least 1152 trials per session. This number of trials resulted from the fully counterbalanced experimental factors: Cued location (6 levels: 1-6 stimuli), initial position of the stimulus array (2 levels: 0, 30 degrees), motion direction (3 levels: CW, CCW, static), presence of continuous intra-saccadic motion (2 levels: absent, present), and delay between the displacement/continuous motion and the masks, i.e., the surface-feature duration (6 levels: 0, 25, 50, 100, 200, 600 ms), thus resulting in a total of 8 trials per experimental cell. Trials were repeated at the end of each session if fixation control was not passed, primary saccades did not reach the pre-saccadic target position, or multiple saccades were made to reachit. On average, observers completed 3705 (SD = 209) trials during all experimental sessions (including repeated and later excluded trials).

Pre-processing involved three major steps. First, 0.5% (SD = 0.4%) of trials were excluded due to unsuccessful fixations (within a central circular boundary of 1.5 dva radius) and dropped frames.

Second, saccades (i.e., primary, secondary, and tertiary saccades in each trial) were detected using the Engbert-Kliegl algorithm (Engbert & Kliegl, 2003; Engbert & Mergenthaler, 2006) with a velocity factor of 10 and a minimal duration of 15 ms. Prior to saccade detection, eye movement data was downsampled to 1000 Hz using bandlimited interpolation. Each trial’s data was padded with its first and last samples and shifted prior to downsampling in order to compensate for the edge effects and delays introduced by low-pass filtering. Sections of missing data due to blinks or tracking problems were expanded by 40 samples on each side and linearly interpolated, but only if those samples were not collected during the relevant trial interval, i.e., from the onset of the saccade cue until 450 ms after the offset of the stimulus motion. Saccade detection was performed on both eyes, but only data collected from the observer’s dominant eye was analyzed, unless the latter was not available due to missing samples, which occurred in 2.4% (SD = 2.1%) of all trials. In order to achieve a conservative criterion for saccade offset (to remove trials in which stimulus motion was not strictly intra-saccadic), we did not consider above-threshold post-saccadic oscillations (if detected within a window of 50 ms after the first below-threshold sample) to be part of the primary saccade.

Third, on average 10.1% (SD = 7.4%) of the remaining trials were excluded because they failed to satisfy the following criteria: (1) No missing data within the relevant trial interval (see above), (2) detection of one single primary saccade that reached the 2-dva area around the pre-saccadic target location (see Procedure), (3) primary saccade metrics compatible with the instructed 10-dva saccade (i.e., amplitude 6 – 15 dva, peak velocity below 600 dva/s, duration below 75 ms), and (4) strictly intra-saccadic stimulus motion (i.e., motion onset after saccade onset and motion offset before saccade offset, regardless of whether continuous motion was present or not), taking into account a deterministic 8.3-ms video delay of the PROPixx projection system (see Schweitzer & Rolfs, 2019).

Ultimately, an average of 3397 (SD = 310) trials per observer entered further analyses. Across observers, stimulus motion was physically displayed 17.8 ms (SD = 0.5) after saccade onset and ended 10.7 ms (SD = 3.1) before saccade offset (both for saccades detected offline and including all system latencies, see also **Figure 2b**). Mean primary saccade amplitude amounted to 9.1 dva (SD = 0.3), mean primary saccade duration to 43.8 ms (SD = 3.0), and mean primary saccade peak velocity to 327.1 dva/s (SD = 34.1).

### Analysis

#### Secondary saccades

On average, secondary saccades were made in 88.2% (SD = 14.1, Mdn = 95.1) of CCW-trials, in 88.6% (SD = 14.2, Mdn = 94.8) of CW-trials, and in 32.0% (SD = 26.9, Mdn = 26.8) of static-trials. Mean secondary saccade rates were slightly reduced by two observers who rarely made secondary saccades despite intra-saccadic displacements, i.e., in 54.5% and 73.3% of trials. Note that overall secondary saccade probability was largely constant across surface-feature duration (0 ms: M = 86.7, SD = 13.0; 25 ms: M = 88.3, SD = 13.6; 50 ms: M = 89.0, SD = 14; 100 ms: M = 88.8, SD = 14.8; 200 ms: M = 88.24, SD = 14.9; 600 ms: M = 89.0, SD = 14.3) and motion conditions (absent: M = 88.4, SD = 14.3; present: M = 88.4, SD = 13.8). To determine whether secondary saccades were made to target or distractor stimuli, we determined whether the offset of the secondary saccade landed within a 3-dva window around the center of either stimulus. In fact, in CCW- and CW-trials 94.5% (SD = 5.3) and 94.7% (SD = 5.1) of secondary saccades landed within these regions. In static-trials, which did not enter further analyses, 55.6% (SD = 30.4) of secondary saccades were re-fixations in the region around the target stimulus. Tertiary saccades, i.e., saccades following secondary saccades, were made in only 8.3% (SD = 4.4) of CCW-trials, 8.3% (SD = 4.5) of CW-trials, and 1.8% (SD = 2.4) of static-trials, and were therefore not further analyzed.

To analyze the proportion of secondary saccades to the pre-saccadic target, we used logistic mixed-effects regression analyses (Bates et al., 2015), specifying observers as intercept-only random effects. The factors of intra-saccadic motion (levels: absent, present) and surface-feature duration (levels: 0, 25, 50, 100, 200, 600 ms) were treatment-coded as ordered factors, so that the condition intra-saccadic motion absent at 0 ms mask SOA constituted the intercept of each model. Secondary saccade latency – defined as the time in milliseconds passed between the offset of the primary saccade and the onset of the secondary saccade – was analyzed using linear mixed-effects regressions applying the same random effects and contrast coding. Confidence intervals for slopes were determined via parametric bootstrapping with 2,000 repetitions each. Along with confidence intervals, p-values were computed via Satterthwaite’s degrees of freedom method. To test the relevance of experimental manipulations, hierarchical model comparisons were performed using the likelihood ratio test and Bayes factors were computed from two models’ respective Bayesian information criteria (Jarosz & Wiley, 2014). While we consistently report the results of model comparisons (pointing out the best model), all reported estimates stem from the full model, not the best model.

To describe the time course of secondary saccade rate to the target, we fitted an exponential growth model with the formula *p*(*t*) = 0.5 + *λ*(1 − *e*^−*β*(*t*−*δ*)^). This model, previously used to describe speed-accuracy tradeoffs (e.g., Carrasco & McElree, 2001), was now used to approximate the proportion of secondary saccades to the target *p*(*t*) – increasing from a chance level of 0.5 – at any given surface-feature duration *t*. We estimated the three parameters of the model, asymptote (λ), slope (β), and onset (δ), in a mixed-effects approach using the stochastic approximation expectation maximization algorithm (starting parameters: λ=1, β=1, δ=4), implemented in the saemix R package (Comets et al., 2017). This approach allowed each of the parameters to be estimated independently for each observer, separately for absent and present intra-saccadic object motion. Subsequently, paired t-tests were used to test whether estimated parameters differed between motion conditions. As we conducted independent hypothesis tests on three parameters, significance levels were Bonferroni-corrected, resulting in *α* = .016. All analyses were implemented in R (R Core Team, 2015) and can be found in a markdown document on OSF: https://osf.io/uafsk/. Furthermore, to describe the time course of secondary saccade latencies, mixed-effects generalized additive models (GAMs) were fitted using the mgcv package in R (Wood, 2017). These models, fitted separately for secondary saccades to the target and to the distractor, allowed to capture the non-linear dynamics of secondary saccades latencies over surface-feature durations, for both experimental conditions of intra-saccadic motion (treatment-coded as ordered factor; reference smooth: absent, difference smooth: present) and for each observer. Thin-plate regression splines (Wood, 2003) were used as smooth functions. **Figure 3** shows the model predictions averaged across observers.

**Figure 3.**
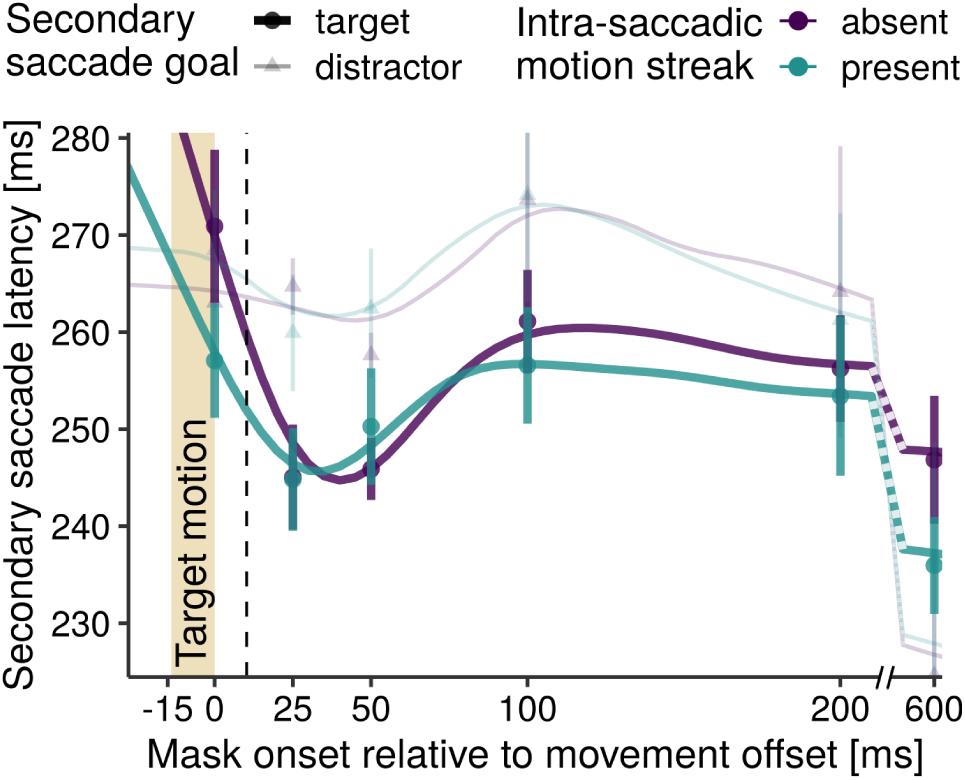
Secondary saccade latency across observers when making secondary saccades to either the initial pre-saccadic target (thick lines and circles) or the distractor (thin, transparent lines and triangles), depending on surface-feature duration and presence of intra-saccadic motion (purple vs green points and lines; error bars indicate ±*S EM*). The beige area indicates the temporal interval of target motion, the vertical dashed line shows the average time of saccade offset after motion offset. Solid lines are predictions of two mixed-effects generalized additive models that describe the time course of observers’ secondary saccade latencies as a function of increasing surface-feature duration. Parametric coefficients of the models indicated an overall significant reduction of secondary saccade latency in the motion-present condition when saccades were directed to the target (Estimate = −5.99, t = −2.21, p = .028), but not when they were directed to the distractor (Estimate = 2.38, t = 0.49, p = .624). The models’ difference smooth terms further suggested a time course modulation due to intra-saccadic motion for target-bound secondary saccades (edf = 9.91, F = 2.99, p = .001), but again not for distractor-bound secondary saccades (edf = 1.01, F = 0.04, p = .836).

#### Reverse Regression

As a first step, target noise patches were convolved with Gabor filters (in sine and cosine phase) of varying orientations (from 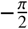 to 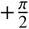,in steps of 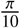 rad) and spatial frequencies (0.25, 0.29, 0.34, 0.39, 0.46, 0.54, 0.63, 0.73, 0.86, 1 cpd), resulting in one energy map per noise patch, that contained the filter responses for each orientation-SF component (an example is shown in **Figure 5a**, see also Schweitzer & Rolfs, 2020; Wyart et al., 2012).

Second, we estimated the angle of the stimulus’ motion trajectory on the retina (for an illustration see **Figure 5b**). In order to compute this retinal trajectory, we subtracted the gaze positions during stimulus presentation (spline-interpolated to match projector refresh rate of 1440 Hz) from the stimulus locations over time. From retinal positions, retinal angles were computed, whose median was subsequently used to normalize each stimulus’ orientation components for its respective retinal trajectory. Relative orientation is the angular difference between the retinal angle and the orientations contained in a given noise patch (Schweitzer & Rolfs, 2020). To achieve the equal-sized steps of relative orientations (in the face of retinal angles that naturally varied between trials), the filter responses for the defined orientation and SF levels were interpolated based on a full tensor product smooth (using cubic splines) of each stimulus’ energy map. Finally, relative orientation could take any value between 0 (orientation parallel to motion direction) and 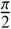 (orientation orthogonal to motion direction).

Finally, we fitted mixed-effects logistic and linear regressions (random intercepts and slopes for observers; Bates et al., 2015) to predict secondary saccades to the target stimulus and inverse latencies of target-bound saccades, respectively, from standardized filter responses in each combination of relative orientation and SF of the target’s energy map. This was done separately for both motion-present and motion-absent conditions (**Figure 5c**,**d**), as well as for surface-feature durations. A significant positive slope for filter responses in a particular relative orientation-SF component indicated that this component drove secondary saccades to the target stimulus or secondary saccades with reduced latency, respectively. Instead of reporting the weights of the model, we reported the corresponding z- and t-statistics, i.e., the ratio of the estimated weights and their standard errors, as they allowed for a more straightforward and comparable evaluation of significance. We further analyzed these outcomes with GAMs. Smooth terms for relative orientation (continuous: 0 .. 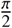) and SF (continuous, log_10_-transformed: −0.6 .. 0), as well as their interactions, were again based on thin-plate regression splines (Wood, 2003) and could include by-variables coding the experimental condition of intra-saccadic motion (treatment-coded as ordered factor; reference smooth: absent, difference smooth: present). Surface-feature duration (reference smooth: 0 ms, difference smooths: 25, 50, 100, 200, 600 ms) was also added in a full model which can be found in the Open Methods (OSF link: https://osf.io/uafsk/). Results shown in **Figure 5c-d** are averages across all surface-feature durations, thus equally taking into account the effect of object features in both movement-present and movement-absent conditions. For each coefficient of the GAM, a complexity of the smooth term (i.e., estimated degrees of freedom, edf) and the significance of the term were estimated. As these estimates cannot be interpreted directly, we complemented the GAM with a simple multiple regression (LM) with the same variable coding to report the linear trends within the data.

## Results

### Efficiency of gaze correction

#### Post-saccadic surface features and intra-saccadic motion drive gaze correction

Of central interest to our research question was whether in case of intra-saccadic target displacement continuous motion would lead to a higher proportion of gaze correction, that is, secondary saccades to the target stimulus. Crucially, intra-saccadic displacements – whether continuous or apparent – had to occur exclusively during saccades, as extending intra-saccadic stimulus manipulations beyond saccade offset has been shown to drastically alter detection performance and subjective appearance of stimuli (Balsdon et al., 2018; Bedell & Yang, 2001; Castet et al., 2002; Duyck et al., 2016; Matin et al., 1972), despite the finding that the window of saccadic suppression often exceeds the saccade duration (Volkmann, 1986). Having excluded those displacements that were not strictly intra-saccadic (see Pre-processing), presentations finished on average 10.7 ms (SD = 3.1) prior to saccade offset (**Figure 2b**) and there was no difference between motion-absent and motion-present conditions (M_absent_ = 10.68, SD_absent_ = 3.08, M_present_ = 10.75, SD_present_ = 3.08; paired t-test: t(9) = 1.24, p = .243).

We first investigated the time course of stimulus feature processing after the stimulus displacements were concluded. We hypothesized that the more time stimulus features were presented before being occluded by masks (**Figure 2a**), the more likely would observers orient their gaze towards the pre-saccadic target, a behavior that likely reflects object correspondence established by comparing post-saccadic object features with those represented in VSTM (Hollingworth et al., 2008; Richard et al., 2008). Indeed, as shown in **Figure 2c**, in the absence of continuous intra-saccadic motion, proportions of secondary saccades made toward the target stimulus increased rapidly and fairly linearly for surface-feature durations of 0 to 50 ms (25 ms: β = 0.53, z = 6.91, 95% CI [0.39, 0.69], p < .001; 50 ms: β = 1.10, z = 13.77, 95% CI [0.94, 1.26], p < .001), reaching an asymptote at 100 ms after displacements occurred (100 ms: β = 1.47, z = 17.51, 95% CI [1.31, 1.65], p < .001; 200 ms: β = 1.45, z = 17.29, 95% CI [1.30, 1.63], p < .001; 600 ms: β = 1.42, z = 17.02, 95% CI [1.27, 1.59], p < .001). Note that group averages reached this asymptote at a comparably low proportion of 80.1% (SD = 5.3). This result was caused by three observers who selected the initial pre-saccadic target on a proportion of trials that was barely above chance (i.e., 55.2%, 51.2%, and 56.9% at the maximum surface-feature duration of 600 ms). There was however no reason or pre-registered criterion for their exclusion.

When the intra-saccadic motion was absent and the surface features were masked right after displacement, the proportion of secondary saccades to the target was at chance level (β_0_ = 0.03, z = 0.12, 95% CI [− 0.39, 0.44], p = .905), as no information was available to perform gaze correction. Crucially, when continuous intra-saccadic motion was present, whereas post-motion surface features were unavailable, secondary saccades were made to the target in 56.1% of trials (SEM = 3.4).This data suggests that gaze correction was accurate significantly above chance (β = 0.25, z = 3.37, 95% CI [0.12, 0.41], p < .001). Although this effect slightly diminished with increasing surface-feature duration (**Figure 2d**), we found no significant interactions for any longer surface-feature duration (25 ms: β = −0.11, z = −1.04, 95% CI [−0.32, 0.10], p = .298; 50 ms: β = −0.06, z = −0.52, 95% CI [−0.29, 0.15], p = .602; 100 ms: β = −0.19, z = −1.61, 95% CI [−0.42, 0.03], p = .106; 200 ms: β = −0.20, z = −1.70, 95% CI [−0.44, 0.03], p = .09; 600 ms: β = −0.04, z = −0.37, 95% CI [−0.29, 0.18], p = .710). Thus, the effect of intra-saccadic motion was largely additive to the effect of surface-feature duration. Hierarchical model comparisons provided further evidence for this view: Adding the factor intra-saccadic motion to a model involving only surface-feature duration (i.e., an additive model) improved the fit to a significant degree (BF_01_ = 1130.04, ΔLL = +11.00, χ^2^(1) = 22.02, p < .001), whereas the full model (including also the interaction term) was not preferred to the more parsimonious additive model (BF_01_ < 0.001, ΔLL = +2.3, χ^2^(5) = 4.61, p = .464).

Taken together, these results suggest that continuous intra-saccadic object motion increased the probability of making a secondary saccade to the cued target. Although the difference between present and absent conditions decreased with increasing surface-feature durations (**Figure 2d**), the performed model comparisons favored a global benefit across surface-feature durations.

#### Intra-saccadic motion results in early onset of information accumulation for gaze correction

What is the nature of the effect of intra-saccadic object motion in the gaze correction paradigm? To find out, we performed an exploratory analysis: We fitted an exponential model (see Analysis) to the probability of making a secondary saccade to the target (**Figure 2c**). Following this procedure, we estimated three parameters of the time course, i.e., asymptote (λ), slope (β), and onset (δ), for motion-absent vs motion-present conditions. We adopted a mixed-effects approach that allowed the three parameters to vary independently for each observer (Comets et al., 2017), so that paired hypothesis tests could be performed. Mean estimates are shown in the table embedded in **Figure 2c**.

Several hypotheses about the benefit of intra-saccadic motion could thus be addressed. First, intra-saccadic motion may result in a gain increase for post-saccadic object features. In this case, it would be expected that performance in the motion-present condition has the same time of onset, but then reaches a higher asymptote. Indeed, estimated λ were slightly larger in the motion-present condition (λ_present_ = 0.304, SE_present_ = 0.058) than in the motion-absent condition (λ_absent_ = 0.291, SE_absent_ = 0.061), but this difference did not reach significance (t(9) = −1.34, p = .214). Second, intra-saccadic motion may lead to an increase of the rate with which post-saccadic information is accumulated, as indicated by the slope of the exponential model. Estimates of β, however, did also not differ between conditions (β_present_ = 0.027, SE_present_ = 0.003, β_absent_ = 0.029, SE_absent_ = 0.003, t(9) = 0.49, p = .634), providing no evidence for such rate increase. Third, despite the fact that all object displacements were finished strictly while the eye was still in flight, continuous intra-saccadic object motion may have revealed the post-saccadic location of the target at an earlier stage, thus allowing the onset of information accumulation to occur already during the ongoing motion. Indeed, estimates of the onset parameter δ revealed a significant difference between the two conditions (δ_present_ = −6.969, SE_present_ = 2.076, δ_absent_ = 0.446, SE_absent_ = 0.053, t(9) = 3.45, p = .007). The results of this analysis suggest that the observed benefit is mainly caused by an earlier availability of object location, which is revealed during intra-saccadic object motion.

#### Post-saccadic surface features and intra-saccadic motion reduce the latency of gaze correction

Given that the presence of intra-saccadic motion increased the likelihood of secondary saccades to the pre-saccadic target in a way consistent with an earlier onset of post-saccadic target localization, we next analyzed the latency of secondary saccades (**Figure 3**). We expected a facilitation of secondary saccade latencies when directed towards the target, but not when directed towards the distractor. On average, secondary saccades to target stimuli were initiated slightly, but insignificantly faster after primary saccade offset (M = 252.3, SD = 37.4; paired t-test: t(9) = 1.35, p = .210) than secondary saccades to distractor stimuli (M = 259.9 ms, SD = 47.6). Average saccade latencies (**Figure 4c**) were well consistent with those found in previous studies using similar paradigms (cf. Hollingworth et al., 2008). We ran linear mixed-effects regression models separately for secondary saccades to the target and distractor to further explore the effects of our experimental design variables onto secondary saccade latency. For secondary saccades made to the target stimulus, and relative to the intercept of the model (no continuous intra-saccadic motion, no surface features), we found significant latency reductions for increasing surface-feature durations (25 ms: β = −26.60, t = −4.87, 95% CI [−37.55, −15.74], p < .001; 50 ms: β = −28.25, t = −5.20, 95% CI [−39.11, −17.60], p < .001; 100 ms: β = −12.89, t = −2.39, 95% CI [−23.07, −2.21], p = .017; 200 ms: β = −19.18, t = −3.54, 95% CI [−30.02, −8.59], p < .001; 600 ms: β = −29.19, t = −5.39, 95% CI [−40.72, −18.31], p < .001), but these latency reductions did not have a linear time course (**Figure 3**). For secondary saccades made to the distractor, we observed no such dependence (25 ms: β = −0.82, t = −0.127, 95% CI [− 13.91, 11.08], p = .898; 50 ms: β = −6.92, t = −1.02, 95% CI [−20.88, 6.72], p = .309; 100 ms: β = 5.87, t = 0.80, 95% CI [−8.40, 19.96], p = .421; 200 ms: β = −3.00, t = −0.40, 95% CI [−18.20, 12.31], p = .690; 600 ms formed an exception: β = −24.75, t = −3.33, 95% CI [−39.45, −10.71], p < .001). Critically, in the absence of surface features, presence of intra-saccadic motion significantly reduced secondary saccade latency to the target (β = −14.27, t = −2.59, 95% CI [−25.43, −3.38], p = .001), but not to the distractor (β = 2.93, t = 0.46, 95% CI [−10.58, 15.41], p = .644). This result was corroborated by a comparison of models including only surface-feature duration and additive models also including the factor intra-saccadic motion: The additive model was slightly better at explaining latencies of secondary saccades to the target (BF_01_ = 1.72, ΔLL = +3.00, χ^2^(1) = 5.6, p = .018), but not to the distractor (BF_01_ = 0.03, ΔLL = +0.01, χ^2^(1) = 0.03, p = .856). Moreover, adding the interaction term improved neither the target model (BF_01_ < 0.001, ΔLL = +4.0, χ^2^(5) = 7.46, p = .189), nor the distractor model (BF_01_ < 0.001, ΔLL = +0.9, χ^2^(5) = 1.81, p = .874), suggesting that the effect of intra-saccadic stimulus motion on secondary saccade latency was additive. In fact, for both target and distractor models, none of the interactions between intra-saccadic motion and suface-feature duration reached significance, except when making secondary saccades to the target at a surface-feature duration of 50 ms (β = 18.01, t = 2.39, 95% CI [3.20, 32.45], p = .017), in which the effect of the motion streak is reversed with respect to the 0 ms condition (see **Figures 3** and **4d**).

**Figure 4.**
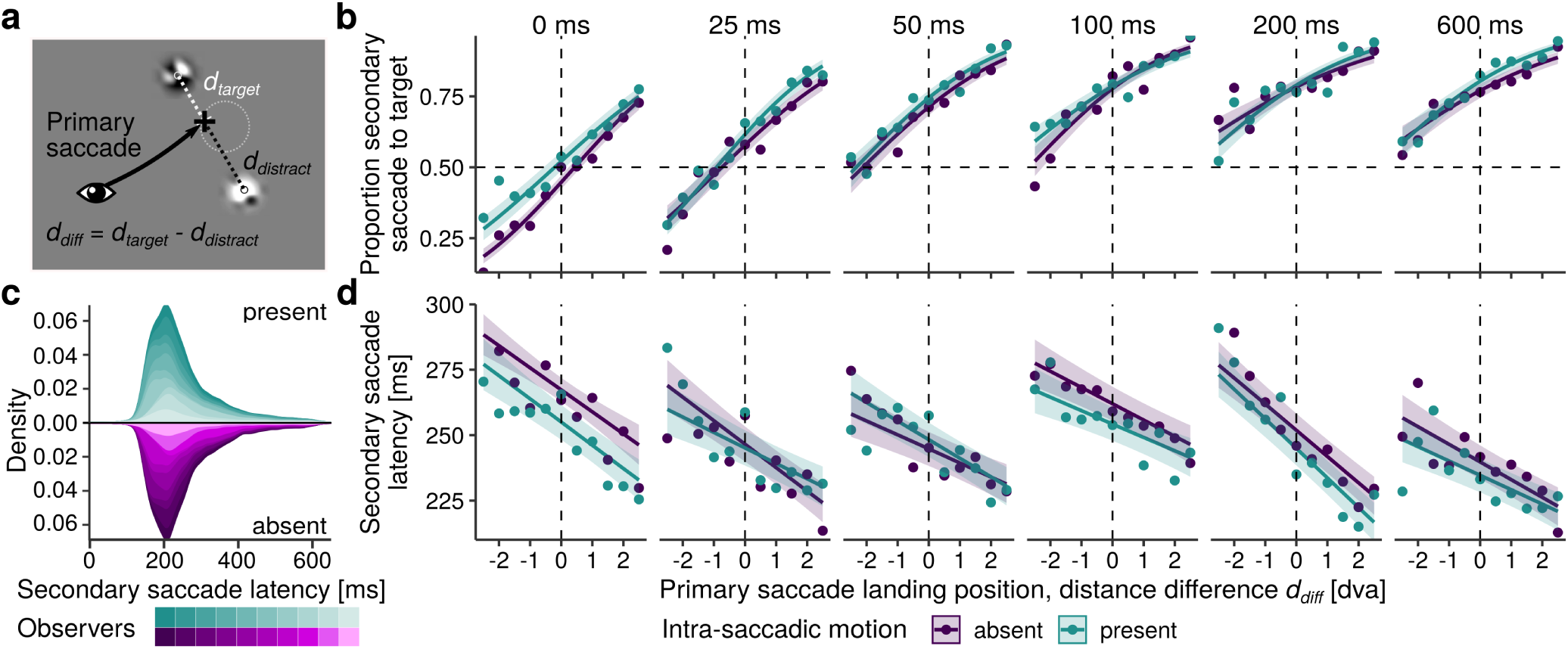
Primary saccade landing positions influence gaze correction. **a** d_diff_ was defined as the difference between the distance from primary saccade landing to the target and the distance to the distractor. Positive values denote that saccades landed closer to the target than to the distractor. **b** Logistic fits modeling the relationship between d_diff_ and the proportion of making a secondary saccade to the target for motion-absent (purple) and motion-present (green) conditions. Panels show results for each surface-feature duration separately. Points indicate group means per 0.5-dva bin. Shaded error bars indicate 95% confidence intervals determined by parametric bootstrapping. **c** Distributions of secondary saccade latencies for each observer. Upper and lower densities represent the motion-absent and motion-present condition, respectively. **d** Linear fits predicting secondary saccade latency to the target stimulus based on d_diff_, surface-feature duration, and presence of intra-saccadic motion.

Taken together, the presence of intra-saccadic stimulus motion thus not only increased the proportion of secondary saccades to the initial pre-saccadic target, but also reduced their latency. A notable detail of the latter result is that the estimated reduction of 14.27 ms (95% CI [25.43, 3.38]) in secondary saccade latency (when no surface features were available) is remarkably similar to the duration of intra-saccadic motion, i.e., 14.6 ms.

### Primary saccade landing positions influence gaze correction

If more than one candidate object for post-saccadic gaze correction is available, a secondary saccade often goes to the closer one (Hollingworth et al., 2008). To investigate a potential interaction of this effect with our observed influence of surface-feature duration and intra-saccadic object motion, we conducted the following analysis. For each trial, we computed the Euclidean distance from the landing position of the primary saccade to the center of the target and to the center of the distractor (**Figure 4a**). Positive values of the difference between these distances (d_diff_) denote landing positions closer to the target than to the distractor. Subsequently, we used d_diff_ in linear mixed-effects regressions to predict saccades to the target as opposed to the distractor (logistic regression; **Figure 4b**) and secondary saccade latency to the target (linear regression; **Figure 4d**).

In predicting secondary saccades to the target, d_diff_ drastically improved the model fit (compared to a model assuming only surface-feature duration) as an additive predictor (BF_01_ > 10^50^, ΔLL = +576.7, χ^2^(1) = 1153.26, p < .001), but only marginally in its interaction with surface-feature duration (BF_01_ < 0.001, ΔLL = +5.2, χ^2^(5) = 10.55, p = .061). When neither surface features nor intra-saccadic motion were available (0 ms, absent condition), landing 1 dva closer to the target increased the probability of making a secondary saccade to the target by a factor of 1.75 (β = 0.56, z = 12.22, 95% CI [0.47, 0.65], p < .001). As shown in **Figure 4b**, this slope decreased slightly when surface features were available upon landing. This decrease was negligible at shorter (25 ms: β = −0.08, z = −1.20, 95% CI [−0.20, 0.05], p = .229; 50 ms: β = −0.07, z = −1.07, 95% CI [−0.21, 0.06], p = .284; 100 ms: β = −0.03, z = −0.47, 95% CI [−0.17, 0.11], p = .636), but significant at longer surface-feature durations (200 ms: β = −0.21, z = −3.01, 95% CI [−0.36, −0.07], p = .003; 600 ms: β = −0.19, z = −2.78, 95% CI [−0.33, −0.06], p = .006). The presence of intra-saccadic target motion significantly increased the probability of secondary saccades to the target when no surface features were available after the displacement (β = 0.34, z = 3.91, 95% CI [0.17, 0.51], p < .001), an effect that was reduced in strength only at 100 ms (β = −0.28, z = −2.16, 95% CI [−0.54, −0.02], p = .031) and 200 ms (β = −0.30, z = −2.28, 95% CI [−0.56, −0.04], p = .023), but not at other surface-feature durations (25 ms: β = −0.16, z = −1.31, 95% CI [−0.40, 0.08], p = .188; 50 ms: β = −0.15, z = −1.22, 95% CI [−0.40, 0.09], p = .223; 600 ms: β = −0.12, z = −0.95, 95% CI [−0.38, 0.13], p = .342). Again, model comparisons suggested that the effects of the two factors were largely additive; including the presence of intra-saccadic motion improved the fit (BF_01_ = 849.46, ΔLL = +12.3, χ^2^(1) = 23.29, p < .001); its interaction with d_diff_ or surface-feature duration did not (BF_01_ < 0.001, ΔLL = +7.3, χ^2^(11) = 16.75, p = .116).

The same analyses were conducted examining linear mixed-effects models predicting secondary saccade latency, provided that these saccades were made to the target. A distribution of all secondary saccade latencies (M_absent_ = 254.6, SD_absent_ = 39.5, M_present_ = 250.4, SD_present_ = 36.4), stacked across observers, is shown in **Figure 4c**. Again, d_diff_ very well predicted saccade latency, but more so as an additive predictor (BF_01_ > 1050, ΔLL = +121.0, χ^2^(1) = 243.85, p < .001) than combined with its interaction with surface-feature duration (BF_01_ < 0.001, ΔLL = +8.1, χ^2^(5) = 14.41, p = .013). When neither surface features nor intra-saccadic motion were available (0 ms, absent condition), landing 1 dva closer to the target reduced secondary saccade latency by 8.4 ms (β = −8.40, t = −4.25, 95% CI [−12.29, −4.53], p < .001). This effect was unaltered by surface-feature duration (25 ms: β = −0.58, t = −0.22, 95% CI [−5.78, 4.59], p = .825; 50 ms: β = 3.02, t = 1.21, 95% CI [−1.97, 7.95], p = .226; 100 ms: β = 2.23, t = 0.89, 95% CI [−2.73, 7.16], p = .372; 200 ms: β = −1.55, t = −0.62, 95% CI [−6.51, 3.33], p = .533; 600 ms: β = 1.61, t = 0.64, 95% CI [−3.24, 6.53], p = .519). When intra-saccadic motion was present, secondary saccade latencies to the target were reduced by 12.4 ms (0 ms: β = −12.36, t = −2.93, 95% CI [−20.71, −4.15], p = .003). This effect was not reduced significantly for any surface feature duration (25 ms: β = 10.79, t = 1.93, 95% CI [−0.15, 21.83], p = .054, 100 ms: β = 4.65, t = 0.89, 95% CI [−5.62, 15.14], p = .374, 200 ms: β = 5.56, t = 1.06, 95% CI [−4.68, 15.93], p = .287, 600 ms: β = 7.25, t = 1.39, 95% CI [−2.95, 17.40], p = .165), except at 50 ms (β = 15.83, t = 2.98, 95% CI [5.26, 26.35], p = .003). Moreover, there was neither an interaction between intra-saccadic motion and d_diff_ (β = −0.45, t = −0.17, 95% CI [−5.78, 4.85], p = .868) nor any higher-level interaction (for full results, see Open Methods at OSF). Even in the grand means, collapsing over any other variables, we found a small, but significant difference between observers’ secondary saccade latencies in the present vs absent conditions (M_absent-present_= 4.17, SEM = 1.68; paired t-test: t(9) = 2.48, p = .035). Indeed, model comparisons revealed that a model including the presence of intra-saccadic motion as an additive factor should be preferred to a model including only d_diff_ and surface-feature duration (BF_01_= 7.76, ΔLL = +6.1, χ^2^(1) = 13.57, p < .001). The full model only marginally improved the fit of the model over the additive model (BF_01_< 0.001, ΔLL = +10.0, χ^2^(11) = 18.67, p = .067).

### Efficient motion streaks facilitate gaze correction

Finally, to establish which stimulus features drive secondary saccades to the target stimulus, we performed a pre-registered, large-scale reverse regression analysis (see Methods). Both contrast sensitivity for moving stimuli and motion perception – especially of high-SF stimuli – are known to dissipate at saccadic velocities (e.g., Burr & Ross, 1982; Castet et al., 2002; Schweitzer & Rolfs, 2019). We hypothesized, therefore, that the rapid movement of the target across the retina produced intra-saccadic motion streaks (Bedell & Yang, 2001; Brooks et al., 1980; Duyck et al., 2016; Geisler, 1999; Matin et al., 1972). In this case, secondary saccades to the target should be facilitated if stimulus features (incidentally) produced a distinctive streak, for instance if the orientation of a stimulus is parallel to its trajectory on the retina (Schweitzer & Rolfs, 2020).

As the noise patches used in this task could potentially contain all possible orientations, as well as SFs from 0.25 to 1 cpd, it was possible to describe each noise patch – both target and distractor – in terms of energy per SF-orientation component (see Methods for details). In brief, for each trial (regardless of whether intra-saccadic target motion was absent or present), we obtained a filter response map for target stimulus by convolving the noise patch with a bank of Gabor filters (**Figure 5a**). Next, we extracted the angle of the target’s trajectory across the retina, which was determined by the target trajectory presented on the screen and the gaze trajectory during presentation (for an illustration, see **Figure 5b**). We then normalized stimulus orientations using this retinal angle, resulting in a measure of relative orientation. As a consequence, stimulus orientations parallel to the retinal angle would result in relative orientations of 0 degrees, whereas stimulus orientations orthogonal to the retinal angle would result in relative orientations of 90 degrees. Finally, we first ran logistic mixed-effects regressions to predict secondary saccades to the target (as opposed to the distractor) from the filter responses present in all available target stimuli. Second, a linear version of the analysis was performed to predict fast saccadic reactions from the same filter responses, provided that secondary saccades were made to the target. Note that a positive relationship between filter responses in a given SF-orientation component and the respective dependent variable implies that this component is either beneficial for gaze correction to the target (**Figure 5c**) or drives secondary saccades to the target at low latencies (**Figure 5d**).

**Figure 5.**
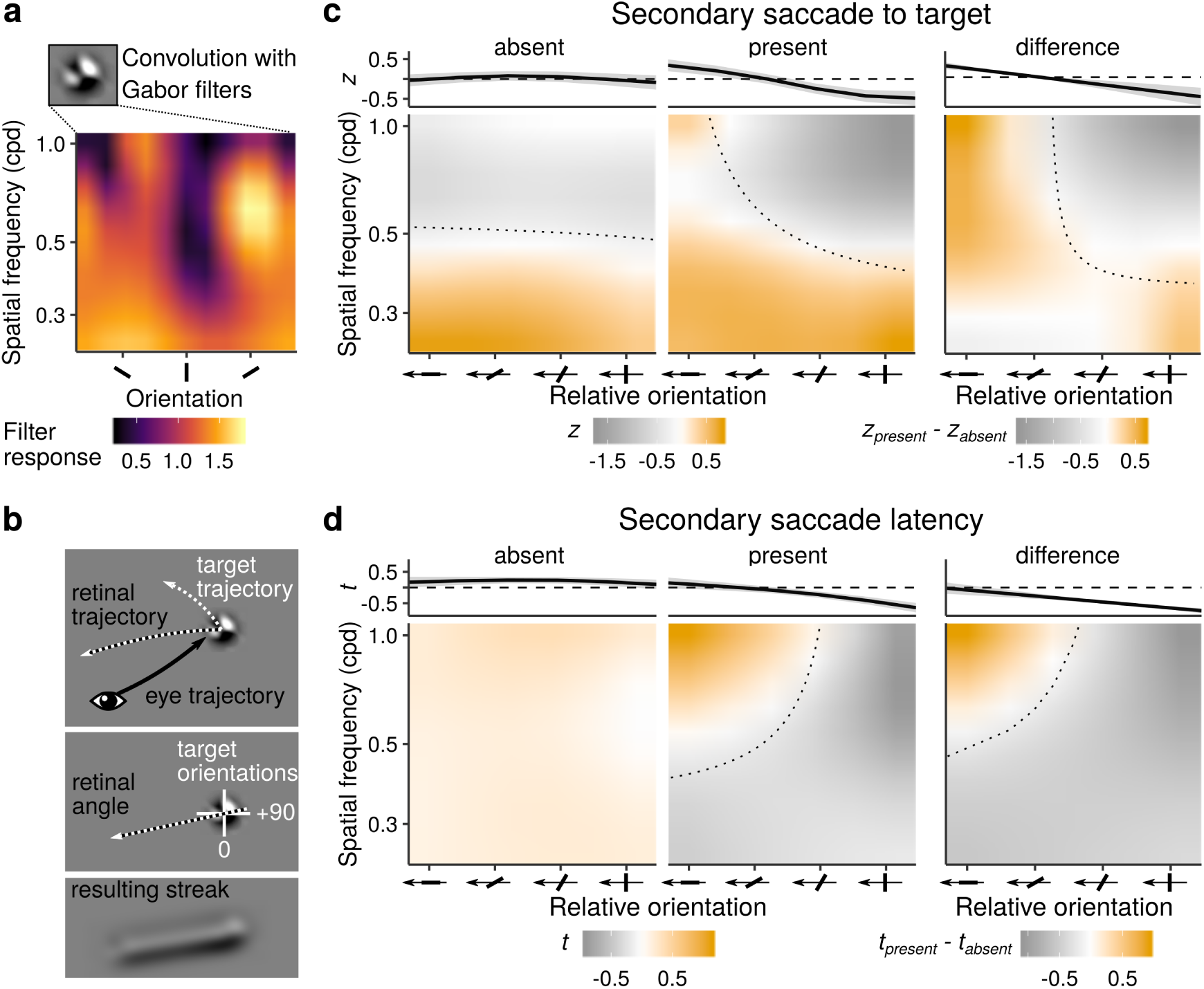
Efficient motion streaks facilitate gaze correction. **a** Example of a filter energy map computed by convolving the noise patch stimulus with a bank of Gabor filters. **b** The retinal trajectory of the target stimulus is the vector sum of the stimulus’ trajectory presented on the screen and the eye position vector during presentation. We computed relative orientation by normalizing the stimulus’ orientation components using the angle of the retinal trajectory. As illustrated by motion filtering applied to the noise patch, orientations parallel to the stimulus’ motion trajectory on the retina should lead to distinctive motion streaks. **c** Results from the logistic reverse regression analysis, fitted by the multivariate GAM, averaged across all surface-feature durations. High z-scores (orange) imply that filter responses in a given SF-orientation component predict the occurrence of a secondary saccade to the target when intra-saccadic motion was present (middle panel) and absent (left panel). Dotted lines demarcate the transition from negative to positive z-scores estimated by the linear model corresponding to the GAM. Upper marginal means show the effect of relative orientation averaged across all SF components. The surface difference (right panel) clearly indicates that secondary saccades to the target (and not to the distractor) were driven by stimulus orientations parallel to the stimulus’ retinal trajectory, suggesting a role of temporal integration of fast-moving stimuli, i.e., motion streaks. **d** Results from the linear reverse regression analysis, using the inverse latency of secondary saccades made to the target as a dependent variable. High t-scores (orange) mark those SF-orientation components facilitating short saccadic reaction times. The pattern suggests that the same parallel orientations that drove secondary saccades to the target also reduced their latencies.

When intra-saccadic motion was absent (**Figure 5c**, left panel), low SFs predicted secondary saccades to the stimulus better than high SFs (GAM: edf = 4.02, F = 15.38, p < .001; LM: β = −2.48, t = −5.13, 95% CI [− 3.43, −1.53], p < .001), whereas relative orientation did not have any impact on gaze correction (GAM: edf = 1.00, F = 0.16, p = .689; LM: β = 0.19, t = 1.02, 95% CI [−0.17, 0.54], p = .306). By contrast, when intra-saccadic motion was present, saccades to target stimuli were driven by smaller relative orientations (GAM: edf = 1.00, F = 15.43, p < .001; LM: β = −1.53, t = −5.97, 95% CI [−2.04, −1.03], p < .001), peaking at relative orientations close to zero (i.e., orientations parallel to the retinal trajectory; **Figure 5c**, middle panel). Moreover, although low SFs were still most relevant, the difference between low and high SFs was reduced (GAM: edf = 2.14, F = 5.01, p = .005; LM: β = 1.43, t = 2.09, 95% CI [0.08, 2.77], p = .036), suggesting that high SFs played a larger role when intra-saccadic motion was available. Crucially, a significant interaction between SFs and relative orientation in the motion-present condition suggested that high SFs were not simply globally more influential, but gained relevance at relative orientations close to zero (GAM: edf = 1.01, F = 21.97, p < .001; LM: β = −3.33, t = −4.63, 95% CI [−4.74, −1.91], p < .001), that is, when (high-SF) stimulus orientations were parallel to the stimulus’ retinal trajectory. This interaction was not present in the movement-absent condition (GAM: edf = 1.0, F = 2.24, p = .134; LM: β = 0.75, t = 1.47, 95% CI [−0.25, 1.74], p = .143). The right panel of **Figure 5c** shows the difference surface of GAM fits for the two experimental conditions. Both conditions were similar with respect to the high predictive value for low-SF components, suggesting that mainly low spatial frequencies served as cues to initiate secondary saccades to target and distractor stimuli. This result seems plausible, not only because post-saccadic stimulus locations were in the visual periphery, but also because filter responses to low SFs were more dissimilar between distractor and target than to high SFs (due to the way luminance was added or subtracted to make the noise patches dissimilar; see Methods), and therefore allowed for better discrimination between the two stimuli. For instance, filter responses to the target and filter responses to the distractor were positively correlated at a SF of 1 cpd (Pearson correlation coefficient, r(19176) = 0.235, 95% CI [0.222, 0.249], p < .001), but strongly negatively correlated at a SF of 0.25 cpd (r(19176) = −0.655, 95% CI [−0.663, −0.647], p < .001). However, low SFs were beneficial in both movtion conditions. Close inspection of the difference surface suggests that mid- and high-SF information drove saccades to the target only when intra-saccadic motion was presented and orientations were close to parallel to the target’s retinal trajectory.

Finally, the analysis of the linear relationships between filter responses and inverse secondary saccade latency revealed that a similar principle applied to the generation of low-latency secondary saccades to the target. In the absence of intra-saccadic motion (**Figure 5d**, left panel), secondary saccade latency was influenced by neither SF (GAM: edf = 1.00, F = 0.26, p = .611; LM: β = 0.23, t = 0.55, 95% CI [−0.59, 1.06], p = .580), nor relative orientation (GAM: edf = 1.81, F = 0.99, p = .369; LM: β = −0.08, t = −0.53, 95% CI [−0.39, 0.23], p = .598), or their interaction (GAM: edf = 1.39, F = 0.39, p = .710; LM: β = −0.14, t = −0.33, 95% CI [−1.02, 0.73], p = .745). In contrast, when motion was present (**Figure 5d**, middle panel), effects of both SF (GAM: edf = 2.43, F = 3.65, p = .012; LM: β = 2.67, t = 4.47, 95% CI [1.50, 3.84], p < .001) and relative orientation (GAM: edf = 1.00, F = 14.55, p < .001; LM: β = −1.21, t = −5.42, 95% CI [−1.66, −0.77], p < .001), as well as a significant interaction between these two predictors (GAM: edf = 1.06, F = 14.91, p < .001; LM: β = −2.52, t = −4.02, 95% CI [−3.75, −1.22], p < .001), were present. This finding suggests that secondary saccade latencies decreased when continuously moving targets had higher energy around SFs of 1 cpd and relative orientations of zero, in other words, when (relatively) high-SF orientations parallel to the target’s retinal trajectory were present. These components were thus able to not only drive secondary saccades to the target, but also enhanced the speed of their initiation.

## Discussion

With each saccade we make, visual objects move rapidly across our retinae, producing transient blurred motion trajectories lawfully related to the ongoing movement. In this study, we emulated these trajectories using a projection system capable of displaying continuous object motion (as opposed to apparent motion from a simple displacement) strictly during saccades with high spatiotemporal fidelity. This technique allowed us to investigate the novel hypothesis that intra-saccadic information about the changing position of saccade targets might facilitate post-saccadic gaze correction to these targets. We tightly controlled the post-saccadic availability of surface features which have been shown to play a crucial role in gaze correction tasks (Hollingworth et al., 2008; Richard et al., 2008), by presenting pixel masks at varying delays. This manipulation allowed the assessment of the impact of intra-saccadic motion on the proportion and latency of secondary saccades to the target, in addition to the time course of the processing of object features.

Even when little or no post-saccadic object information was available, the presence of intra-saccadic target motion increased the rate of secondary saccades to the original pre-saccadic target and reduced their initiation latency. These results are central to our hypothesis, as they suggest that intra-saccadic information was not suppressed or otherwise omitted – as widely assumed (for a review, see Castet, 2010) – but instrumental for timely gaze correction.

The magnitudes of these effects may seem small at first, but they were consistent with what was to be expected from a 14.6-ms intra-saccadic motion duration: Information about post-displacement object features was accumulated in a exponential fashion right upon motion onset. A comparison of the parameters of these exponential functions suggests that facilitation caused by intra-saccadic motion was not due to an increase of gain or acculumation rate when processing object features and locations, but due to the earlier availability of these, starting with the onset of intra-saccadic target motion. Notably, even when continuous object motion was absent, the models predicted that at saccade offset (on average 10 ms after object motion offset) secondary saccade rates to the target would already be above chance, suggesting that visual processing starts before saccade landing. Consistent with this view, the estimated secondary saccade latency reduction was again of the same magnitude as the motion duration both when surface features were unavailable and when they were available for the entire 600 ms. Note that, in this task, the motion duration was barely a third of the mean saccade duration. In natural vision, any visual object could produce motion smear across the entire duration of the saccade, possibly supporting short-latency corrective saccades upon saccade landing.

Furthermore, we not only showed that the effect of intra-saccadic object motion is orthogonal to the effect of primary saccade landing positions (cf. Hollingworth et al., 2008), but also provided evidence for the benefit of effective temporal integration when stimulus orientations were aligned with their retinal motion trajectories – a typical signature of motion streaks (Schweitzer & Rolfs, 2020). In other words, the more effectively the combined movement of eye and target in a given trial generated a motion streak, the more often did a secondary saccade go to the target. The same effect was found for the latency of these saccades, providing evidence that efficient motion streaks provided a benefit not only for the accuracy but also for the speed of gaze correction. Although it has been shown that motion perception during saccades is well possible (Castet et al., 2002), contrast sensitivity to gratings orthogonally oriented to their motion trajectories is drastically reduced at saccadic velocities (Burr & Ross, 1982; Mitrani & Yakimoff, 1970). In contrast, motion streaks often remain well resolved even at saccadic speeds (Bedell & Yang, 2001; Brooks et al., 1980; Duyck et al., 2016; Matin et al., 1972), partly due to visual persistence. This invites the intruiging hypothesis that they might be able to link objects across saccades via spatiotemporal continuity. Our results show that even when objects were displaced while the eyes were in mid-flight, a continuous presence of the target throughout the saccade – as opposed to a very brief disruption of this continuity – facilitated gaze correction, regardless of how long feature information was available after the displacement. Although this facilitation was clearly strongest shortly after motion offset, interactions between surface-feature duration and presence of intra-saccadic motion were rarely significant and all model comparisons performed on both secondary saccade rate and latency data favoured the additive model over the full model. These consistent results suggest that post-saccadic object features and spatiotemporal continuity – established by intra-saccadic continuous object motion – contributed rather independently to gaze correction performance. This conclusion is well in line with the predictions of the object-file theory (Kahneman et al., 1992; Mitroff & Alvarez, 2007), which suggests that objects are bound to spatial indexes. It is also consistent with the view that surface features are functional for the establishment of object correspondence (Hollingworth et al., 2008; Hollingworth & Franconeri, 2009; Richard et al., 2008): Intra-saccadic motion streaks may not only be indicators of amplitude and direction of continuous shifts of objects across saccades, but to some extent also maintain the object’s surface features, such as color, which has been shown to be largely unaltered by saccadic suppression (Burr et al., 1994; Bridgeman & Macknik, 1995; Knöll et al., 2011), throughout the saccade.

To conclude, our results support the idea that saccades do not cause gaps in visual processing, as even motion smear induced by high-velocity, brief, unpredictable, and strictly intra-saccadic object motion was taken into account when performing gaze correction. Depending on the efficiency of intra-saccadic vision especially in real-world visual environments, the visual consequences induced by our very own saccades may constitute an unexpected contribution to achieving object continuity and, through it, visual stability.

## Acknowledgments

R.S. was supported by the Studienstiftung des deutschen Volkes and the Berlin School of Mind and Brain. M.R. was supported by the Deutsche Forschungsgemeinschaft (DFG, grants RO3579/8-1 and RO3579/10-1).

We thank Emilia Maria Rehse for her indispensable work in data collection, as well as Lisa Kröll, Greta Häberle, Frederik Geweke, and Bryce Yahn for their support in the initial stages of the project.

## Author contributions

R.S. and M.R. conceived, designed, and pre-registered the study. R.S. implemented and conducted the experiment. R.S. analyzed data under M.R.’s supervision. R.S. drafted the manuscript, and M.R. provided critical revisions.

## Notes

### Competing Interest Statement

The authors have declared no competing interest.

### Summary of Updates

Figure 5 revised: A second type of reverse regression (using inverse saccade latency as dependent variable) was added.

